# Emergence of a dynamical state of coherent bursting with power-law distributed avalanches from collective stochastic dynamics of adaptive neurons

**DOI:** 10.1101/2024.05.28.596196

**Authors:** Lik-Chun Chan, Tsz-Fung Kok, Emily S.C. Ching

## Abstract

Spontaneous brain activity in the absence of external stimuli is not random but contains complex dynamical structures such as neuronal avalanches with power-law duration and size distributions. These experimental observations have been interpreted as supporting evidence for the hypothesis that the brain is operating at criticality and attracted much attention. Here, we show that an entire state of coherent bursting, with power-law distributed avalanches and features as observed in experiments, emerges in networks of adaptive neurons with stochastic input when excitation is sufficiently strong and balanced by adaptation. We demonstrate that these power-law distributed avalanches are direct consequences of stochasticity and the oscillatory population firing rate arising from coherent bursting, which in turn is the result of the balance between excitation and adaptation, and criticality does not play a role.

## I. INTRODUCTION

Neurons in the brain fire even when they do not receive any external stimuli and such firing activities are known as spontaneous activity (see reviews [1, 2] and references therein). Spontaneous activity is far from random but has rich dynamical structures in space and time. There are low frequency oscillations in the range of 0.01 - 0.2 Hz [3, 4] and collective bursting events, called neuronal avalanches, at time scales shorter than about 100 ms, have been revealed from analyses of in vitro measurements from cortical slices and dissociated cultures of cortical and hippocampal neurons of rats [5–7] and in vivo measurements from anesthetized rats [8] and awake monkeys [9]. These neuronal avalanches have power-law distributions in duration *T* and size *S*:

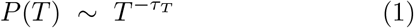

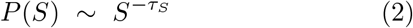

Further analysis [10] shows that the mean temporal profiles of avalanches of varying durations can be described by a scaling function such that the average size of avalanches of given duration *T* has a power-law dependence on *T* :

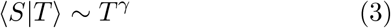

and the exponents obey approximately a scaling relation

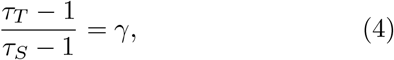

This scaling relation has been derived for the critical phenomenon of crackling noise [11, 12].

The observed power-law distributed neuronal avalanches have attracted particular attention since they have been interpreted as evidence supporting the hypothesis that the brain is operating near a critical point of transition between two states [13–17] and many models of criticality have been proposed for studying brain dynamics [18–34]. However, the values of *τ*_*T*_ and *τ*_*S*_ were found to depend on the time scale of analysis, which is the width Δ*t* of the time bins used in the analysis of the avalanches [5, 9], in contrast to what one expects for a critical system. With Δ*t* taken as the average interevent interval IEI_ave_ of the whole neuronal population [5], further dependence of the exponents on the spiking variability level of in vivo measurements is found and that at some spiking variability level, Eq. (4) approximately holds with (*τ*_*T*_ − 1)*/*(*τ*_*S*_ − 1) ≈1.28 while *τ*_*T*_ and *τ*_*S*_ are different for different animal species [35]. In [36, 37], it has been shown that variations of *τ*_*T*_ and *τ*_*S*_, and Eq. (4) with (*τ*_*T*_ − 1)*/*(*τ*_*S*_ −1) ranging from 1.37 to 1.66 could be obtained using a cortical branching model for quasicritical brain dynamics. This model contains only excitation and lacks biological details [38]. In addition, power-law distributions can be generated by mechanisms other than criticality (see review [39] and references therein) and power-law distributed neuronal avalanches have been shown to exist in stochastic noncritical systems [40–46]. Two of these studies showed that an inhomogeneous Poisson process with a time varying rate can give rise to avalanches of power-law distributions with cutoffs but how such a time varying rate could arise is unclear [43, 45]. Furthermore, most of the previous studies focus on power-law distributed neuronal avalanches and although other studies show that power-law distributed neuronal avalanches and oscillations can co-exist [23, 30–34], it has yet to be shown that oscillations and power-law distributed avalanches with all the features observed in experiments can exist in one model system. Hence, debates on whether or not the brain is operating at criticality are ongoing [47–49] and the origin and underlying mechanism of the observed oscillations and power-law distributed neuronal avalanches in spontaneous brain activity remain to be understood.

In this paper, we study the dynamics of networks of adaptive neurons, subjected to stochastic input, using biologically plausible models of neurons and synapses. We map out the possible dynamics in the parameter space of excitatory and inhibitory synaptic strengths and show the emergence of a dynamical state of coherent bursting with power-law distributed avalanches in a regime of intermediate excitatory synaptic strengths. In this state, each neuron fire in bursts and the bursting dynamics of the neurons are coherent such that the time-binned population firing rate is oscillatory on long time scales. At shorter time scales, the bursting activity forms avalanches that have power-law duration and size distributions satisfying Eqs. (1)-(4) with (*τ*_*T*_ −1)*/*(*τ*_*S*_ −1) ≈1.3 as observed in experiments [35]. We further show that coherent bursting is a result of the balance between excitation and adaption and establish that power-law neuronal avalanches do not only co-exist with the oscillations arising from coherent bursting but they are the consequences of stochasticity and such oscillations, and criticality does not play a role.

## II. MODEL

We study a network of *N* neurons subjected to stochastic input using biologically plausible Izhikevich’s neuron model [50] and conductance-based synapse model [51– 53]. In Izhikevich’s model [50], the dynamics of a neuron is described by a membrane potential *v*_*i*_ and a membrane recovery variable *u*_*i*_, where the label *i* runs from 1 to *N* in the network. The recovery variable *u*_*i*_ accounts for the activation of potassium ionic currents and inactivation of sodium ionic currents, and provides a negative feedback to *v*_*i*_. Neuron *i* receives a synaptic current 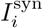 from all its pre-synaptic neurons and a stochastic current *I*_noise_. The time evolution of *v*_*i*_ and *u*_*i*_ is governed by two coupled nonlinear differential equations

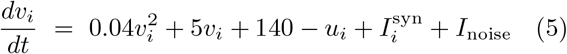

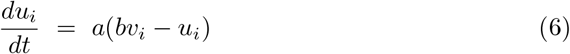

where *v*_*i*_ and *u*_*i*_ are in mV, *t* is time in ms and *a* and *b* are positive parameters. The stochastic current *I*_noise_ = *αξ*, where *α >* 0 and *ξ* is a Gaussian white noise with zero mean and unit variance:

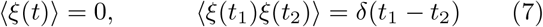

Every time when *v*_*i*_ ≥ 30, neuron *i* fires and a spike is generated, then *v*_*i*_ and *u*_*i*_ are reset:

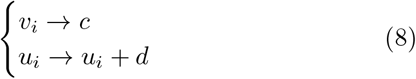

where *c >* 0 and *d* ≥ 0 are parameters. When *d >* 0, *u*_*i*_ and thus the negative feedback increases every time a neuron spikes and this models spike frequency adaptation, which is a reduction in the spiking frequency over time in response to a prolonged stimulus, a property observed in biological neurons. It has been shown that [54] with suitable choices of the parameters *a, b, c*, and *d*, the Izhikevich model is capable of reproducing the rich firing patterns exhibited by real cortical neurons from different electrophysiological classes [55, 56]. We consider only two types of neurons, excitatory regular spiking neurons (*a* = 0.02 and *d* = 8) and inhibitory fast spiking neurons (*a* = 0.1 and *d* = 2) in the network and both types have *b* = 0.2 and *c* = − 65 mV [50].

In a conductance-based synapse model [51–53], the synaptic current is related to the membrane potential:

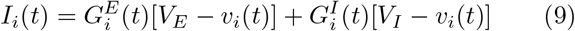

where *V*_*E*_ = 0 and *V*_*I*_ = −80, both in mV, are the reversal potentials of excitatory and inhibitory synapses, and 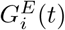 and 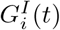 are the excitatory and inhibitory conductances. The conductance 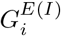 increases by the magnitude of the synaptic weight *w*_*ij*_ whenever a presynaptic excitatory (inhibitory) neuron *j* fires and generates a spike at *t*_*j,k*_, otherwise it decays with a time constant *τ*_*E*(*I*)_ [57–59]:

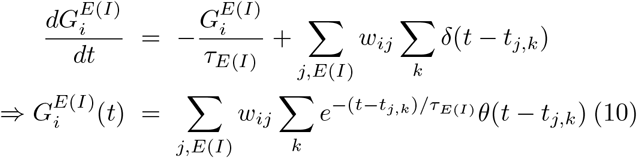

where *τ*_*E*_ = 5 ms and *τ*_*I*_ = 6 ms [60, 61], Σ_*j,E*(*I*)_ sums over all presynaptic excitatory (inhibitory) neurons *j*, and *θ*(*t* − *t*_0_) is the Heaviside step function.

To facilitate theoretical analysis, we focus on a simple directed random network (network A) with the same excitatory and inhibitory incoming degrees (*k*_*E*_ = 8 and *k*_*I*_ = 2) for each neuron and constant excitatory and inhibitory synaptic strength, i.e., *w*_*ij*_ = *g*_*E*_ or *g*_*I*_ for neuron *j* being excitatory or inhibitory. Network A has *N* = 1000 neurons with *N*_*E*_ = 800 excitatory and *N*_*I*_ = 200 inhibitory neurons. To test the robustness of our results, we study two additional networks: network B has the same network structure as network A and the excitatory and inhibitory synaptic weights are taken from uniform distributions with mean values of *g*_*E*_ and *g*_*I*_, and network C is a directed network reconstructed from multi-electrode array data measured from a neuronal culture [62], using the method of directed network reconstruction proposed by Ching and Tam [63], with constant excitatory and inhibitory synaptic strengths *g*_*E*_ and *g*_*I*_. The degree distributions of network C are qualitatively different from those of network A. Network A is homogeneous in the incoming degree by construction while network C is heterogeneous with a bimodal incoming degree distribution and the outgoing degree distribution is long-tailed for network C and binomial for the random network A (see Appendix). Results reported below are for network A unless otherwise stated. Similar results are found in networks B and C and will be discussed in the Appendix.

## III. NUMERICAL RESULTS

The stochastic differential equations (5) and (6) together with equations (9) and (10) were integrated using the weak second-order Runge-Kutta method [64] with a time step *dt* = 0.001 ms. We set the initial values of *v*_*i*_ and *u*_*i*_ to be −70 and −14, respectively and study the dynamics as a function of *g*_*E*_ and *g*_*I*_ at a fixed value of *α* = 3.

### A. Three distinct dynamical states

The dynamics depend on both *g*_*E*_ and *g*_*I*_ and three distinct dynamical states are found. At a fixed value of *g*_*I*_ including *g*_*I*_ = 0, the dynamics changes from state I of irregular and independent spiking to state II of coherent bursting and finally to state III of incoherent fast spiking as *g*_*E*_ increases. The phase diagram of the different dynamical states is presented in Fig. 1. A similar phase diagram is found for network B (see Appendix). Previous studies suggested that the possible behaviors of a network of excitatory and inhibitory neurons can be determined by the ratio *g*_*I*_ */g*_*E*_ among other factors [59, 65] but our results clearly show that the system can be in different dynamical states even when *g*_*I*_ */g*_*E*_ is the same.

**FIG. 1:**
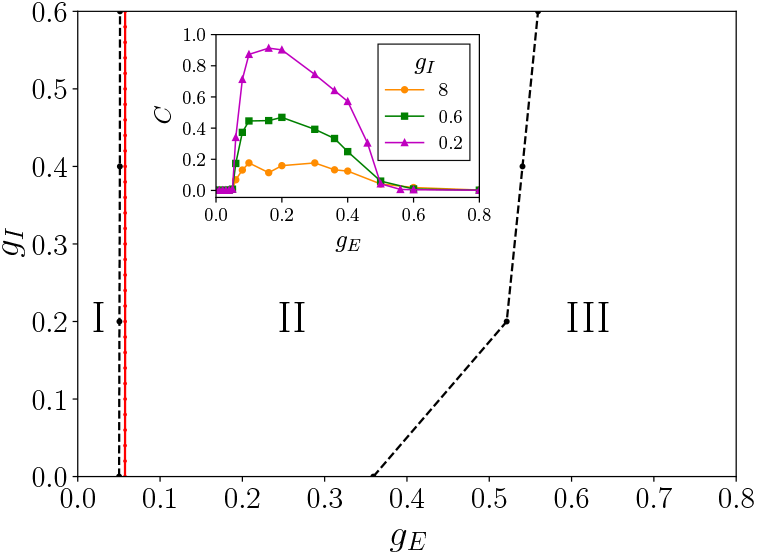
Different dynamical states in the *gE* -*gI* parameter space at α = 3. The solid red line is the theoretical threshold 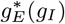 above which state I becomes unstable (see Sec. IV A). In the inset, we show the dependence of the coherence param-eter C on *gE* at given values of *gI* .

In state I, *g*_*E*_ is small such that the synaptic current *I*^syn^ is too weak to initiate a spike on its own. Spiking activity is essentially induced by the stochastic current *I*_noise_. The raster plot, which shows the times at which the spikes occur, thus consists of scattered random dots (see Fig. 2a). In state II, neurons fire in bursts displaying relatively fast spiking that are separated by intervals of quiescence. There is a high coherence in the bursting dynamics of the neurons across the whole network while spikes within a burst from individual neurons are not synchronized. This coherent bursting is a collective effect since an isolated excitatory or inhibitory neuron, subjected to the stochastic input (or a constant input), does not exhibit bursting. The raster plot then consists of stripes of densely distributed dots, corresponding to the bursts (see Fig. 2b). In state III, neurons exhibit fast spiking and some might still fire in bursts but the spiking or bursting is incoherent across the whole network and the raster plot consists of densely distributed dots that spread in time (see Fig. 2c).

**FIG. 2:**
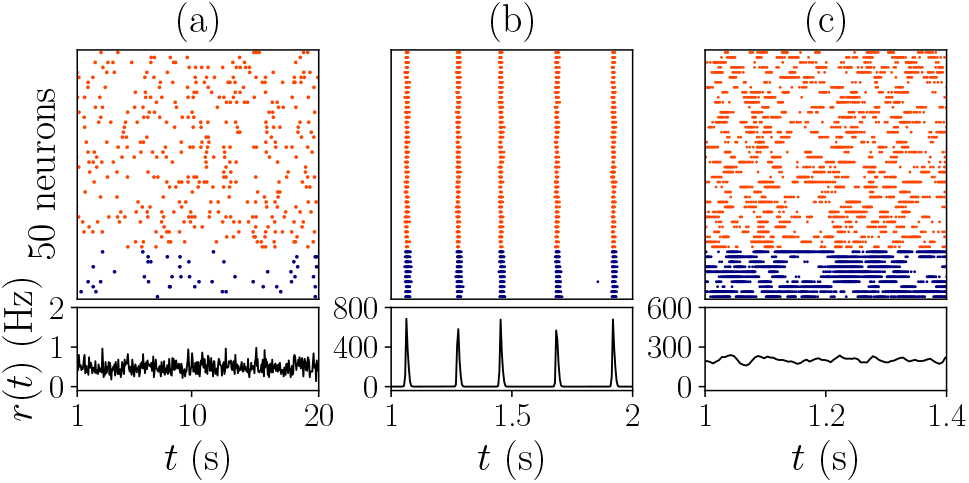
Raster plots for 10 inhibitory (blue) and 40 excitatory (red) neurons randomly chosen and the population averaged time-binned firing rate *r*(*t*) for (a) (*gE*, *gI*) = (0.04, 0.2) in state I, (b) (*gE*, *gI*) = (0.2, 0.2) in state II, and (c) (*gE*, *gI*) = (0.6, 0.2) in state III. The width of the time bin used in calculating *r*(*t*) is 50 ms in (a) and 5 ms in (b) and (c).

Denote the time-binned firing rate of each neuron by *r*_*i*_(*t*), *i* = 1, 2, …, *N* and *t* = *nw, n* = 1, 2, …, *N*_*bin*_, where *w* is the width of the time bin and *N*_*bin*_*w* is the observation time. The population averaged time-binned firing rate is 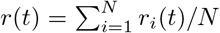. The extent of temporal fluctuation of the spiking activity of each neuron is characterized by the variance of *r*_*i*_(*t*):

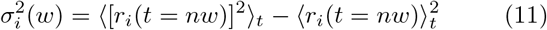

where 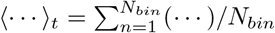. Similarly, the extent of temporal fluctuation of the population averaged spiking activity is characterized by the variance of *r*(*t*):

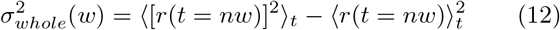

To measure the degree of coherence or synchrony of the neuron dynamics across the network, we define a coherence parameter *C*

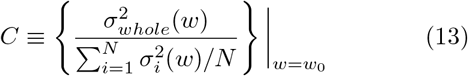

evaluated at *w*_0_ = 32 ms, which is chosen to be about the average duration of the bursts (average width of the stripes in the raster plot) in state II. 0 ≤ *C* ≤ 1 and for the case of total coherence with *r*(*t*) = *r*_*i*_(*t*), *C* = 1. Our definition of the coherence parameter modifies an earlier definition [66] by replacing the membrane potentials of neurons by their time-binned firing rates for the ease of computation. At a given value of *g*_*I*_, *C* increases from about 10^*−*3^ in state I to around 0.1-1 in state II and then decreases to about 10^*−*3^ in state III as shown in the inset of Fig. 1. At a fixed value of *g*_*E*_ in state II, *C* decreases when *g*_*I*_ is increased as the fraction of neurons participating in the network bursts decreases. The fraction of participation also varies from burst to burst leading to a variation in heights, widths and shapes of the peaks in the oscillatory population averaged time-binned firing rate *r*(*t*) (see Fig. 3). We use a threshold value of 0.03 to mark the boundaries between the states in Fig. 1.

**FIG. 3:**
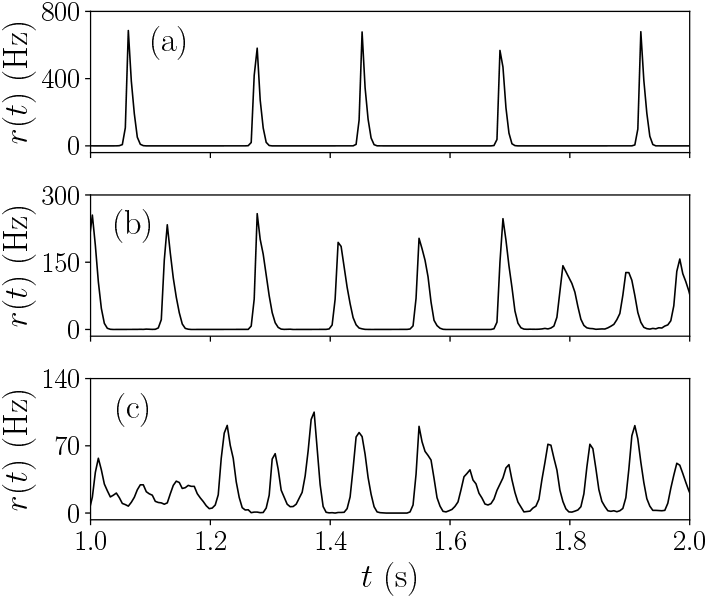
Population averaged time-binned firing rate *r*(*t*) for (a) (*gE*, *gI*) = (0.2, 0.2), (b) (*gE*, *gI*) = (0.2, 0.6), and (c) (*gE*, *gI*) = (0.2, 8) in state II. The width of the time bin used in calculating *r*(*t*)is 5 ms.

### B. Avalanches

Neuronal avalanches refer to cascades of events in which the activities of individual neurons trigger further firing of their postsynaptic neurons. In practice, avalanches are often extracted from measured signals by an operational definition put forward by Beggs and Plenz [5]. In a similar manner, we use the spiking activities of the neurons to define avalanches. The time series of the spiking activities of the whole network are partitioned into time bins with a width of Δ*t*. The time-binned spiking activity of the whole network can be treated as the time-binned activity of an effective neuron, which spikes in a certain time bin whenever any individual neuron spikes in that time bin. A null event of the effective neuron is identified in a certain time bin if there are no spiking activities from all neuron in that time bin. An avalanche is defined as a sequence of spiking activities in consecutive time bins between two null events of the effective neuron. The duration of an avalanche is *T* = *n*Δ*t* where *n* is the number of time bins in its sequence of activity and the size *S* of an avalanche is the total number of spikes of all neurons occurring in its duration.

The distributions of duration and size of the avalanches, *P* (*T*) and *P* (*S*) have similar forms in states I and III. As can be seen in Figs. 4 and 5, *P* (*T*) in these two states are well described by an exponential distribution

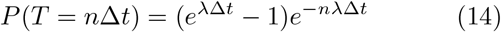

and *λ* decreases when Δ*t* increases. The solid lines in Figs. 3 and 4 are theoretical results to be discussed in Sec IV. Apparently, *P* (*S*) can also be described by an exponential distribution but as we will discuss in Sec. IV, they are only close to an exponential distribution.

**FIG. 4:**
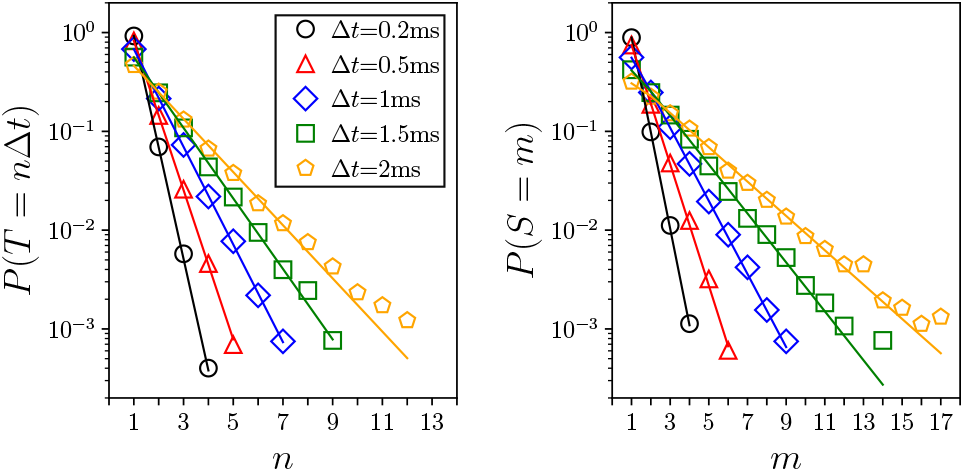
*P*(*T*) and *P*(*S*) for *gE* = 0.03 and *gI* = 0.2 in state I. The solid lines are theoretical results (see Sec. IV).

**FIG. 5:**
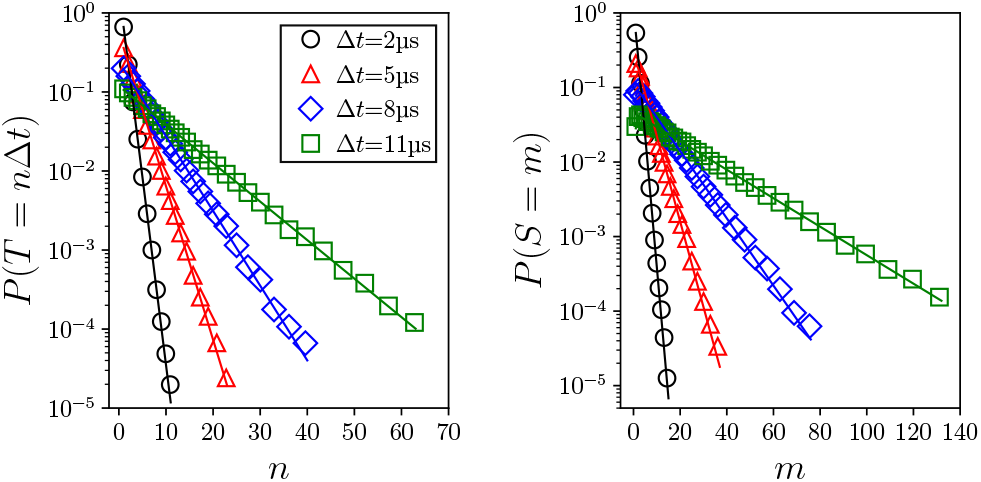
Similar plots as in Fig. 4 for g*E* = 0.6 and g*I* = 0.2 in state III.

In state II, at fixed values of *g*_*E*_ and *g*_*I*_, *P* (*T*) and *P* (*S*) are well described by power laws, and ⟨*S* |*T*⟩ has a power-law dependence on *T* for a range of Δ*t*. The exponents *τ*_*T*_ and *τ*_*S*_ vary with Δ*t* as found in experiments [5, 9] but *γ* remains approximately the same (see Table 1). The power-law exponents for *P* (*T*) and *P* (*S*) are estimated by the maximum likelihood estimator of a discrete power law distribution and the hypothesis that the data are described by a power law is validated by the goodness-of-fit test using the Kolmogorov-Smirnov statistic (with a requirement of *p*-value greater than 0.1) [67]. A common choice of Δ*t* is the average interevent interval, IEI_ave_, of the effective neuron [5]. We find that IEI_ave_ does not always fall within the range of Δ*t* for which power-law duration and size distributions are observed and in these cases we choose a Δ*t* that is around the middle of the range. In Fig. 6, we show *P* (*T*), *P* (*S*) and ⟨*S* |*T*⟩. The results for *τ*_*T*_, *τ*_*S*_ and *γ* for all the cases studied are presented in Table 1. While *τ*_*T*_ and *τ*_*S*_ depend on *g*_*E*_ and *g*_*I*_ or Δ*t*, the relation Eq. (4) holds approximately throughout state II with (*τ*_*T*_ −1)*/*(*τ*_*S*_ −1) ≈ 1.3 as observed in experiments [35] (see Fig. 7). Similar results are found in state II for a range of intermediate values of *g*_*E*_ in network C (see Appendix).

**TABLE I:** The exponents τ*T*, τ*S* and γ for the different cases studied. When Δ*t* ≠ IEI_ave_, the values of IEI_ave_ are indicated in parentheses. The dependence of the exponents on Δt is illustrated for *gE* = 0.3 and *gI* = 0.2.

| $g_E$ | $g_I$ | $\tau_T$ | $\tau_S$ | $\frac{(\tau_T - 1)}{(\tau_S - 1)}$ | $\gamma$ | $\Delta t$ (ms) |
| --- | --- | --- | --- | --- | --- | --- |
| 0.075 | 0 | 2.12 | 1.86 | 1.30 | 1.30 | 0.03 |
| 0.1 | 0 | 1.94 | 1.72 | 1.31 | 1.28 | 0.005 (0.021) |
| 0.2 | 0 | 2.16 | 1.87 | 1.33 | 1.23 | 0.014 |
| 0.075 | 0.2 | 2.04 | 1.79 | 1.32 | 1.29 | 0.04 (0.147) |
| 0.1 | 0.2 | 1.91 | 1.73 | 1.25 | 1.27 | 0.02 (0.066) |
| 0.2 | 0.2 | 1.99 | 1.76 | 1.30 | 1.29 | 0.015 (0.037) |
| 0.3 | 0.2 | 2.08 | 1.80 | 1.35 | 1.30 | 0.027 |
| 0.3 | 0.2 | 1.89 | 1.65 | 1.37 | 1.27 | 0.04 (0.027) |
| 0.3 | 0.2 | 1.92 | 1.68 | 1.35 | 1.27 | 0.05 (0.027) |
| 0.3 | 0.2 | 1.97 | 1.72 | 1.35 | 1.27 | 0.06 (0.027) |
| 0.4 | 0.2 | 1.87 | 1.68 | 1.28 | 1.27 | 0.019 |
| 0.075 | 0.6 | 2.12 | 1.90 | 1.24 | 1.29 | 0.1 (0.195) |
| 0.1 | 0.6 | 2.31 | 2.00 | 1.31 | 1.28 | 0.115 |
| 0.2 | 0.6 | 2.05 | 1.80 | 1.31 | 1.29 | 0.036 |
| 0.3 | 0.6 | 1.92 | 1.70 | 1.31 | 1.28 | 0.02 |
| 0.1 | 8 | 2.13 | 1.90 | 1.26 | 1.31 | 0.16 (0.139) |
| 0.2 | 8 | 2.16 | 1.89 | 1.30 | 1.25 | 0.06 (0.043) |
| 0.3 | 8 | 1.94 | 1.72 | 1.31 | 1.29 | 0.04 (0.02) |

**FIG. 6:**
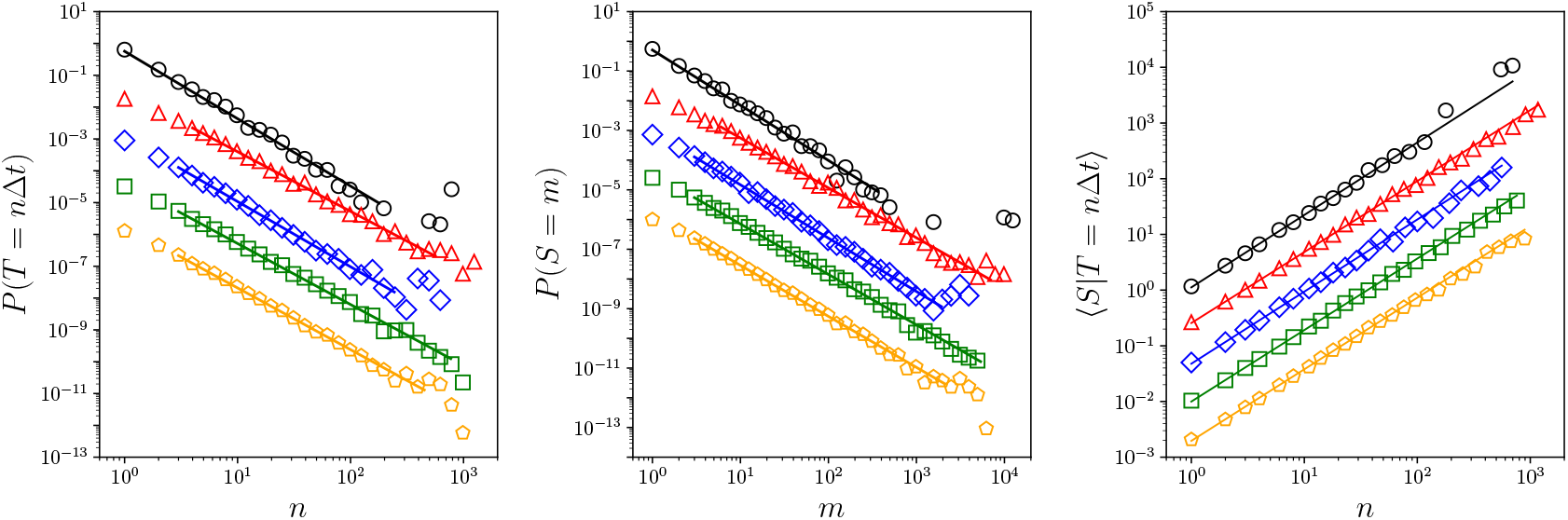
*P*(*T*), *P*(*S*) and ⟨*S*|*T*⟩ for (*g*_*E*_.*g*_*I*_) = (0.075,0), (0.4,0.2), (0.2,0.6), (0.3,0.6) and (0.3,8), from top to bottom, in state II for Δ*t* = IEIave except for (0.3,8) where Δ*t* = 2IEI_aVe_. The curves, other than the topmost ones, are shifted downwards for clarity.

**FIG. 7:**
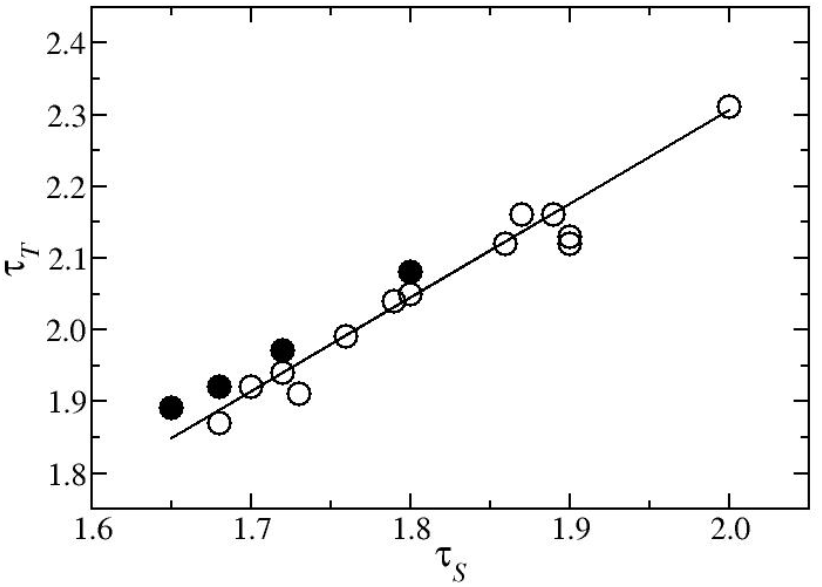
Relation between τ_*T*_ and τ_*S*_ for all the cases in state II studied as shown in Table 1. The filled symbols correspond to (*g*_*E*_.*g*_*I*_) = (0.3,0.2) with different values of *Δt*. The solid line is the least-square fit of (*τ*_*T*_ — 1)/( *τ*_*S*_ — 1) = *K* and the fitted value of *K* is 1.3.

## IV. THEORETICAL ANALYSES AND RESULTS

### A. State I

In state I, neurons fire irregularly, triggered essentially by *I*_noise_, and each neuron approximately fires independently of one another. Thus we approximate the dynamics of state I as *N* independent Poisson processes with event rates equal to the firing rates of the neurons. With this approximation, we can derive *P* (*T*), *P* (*S* | *T*) and *P* (*S*). The time-binned spiking activity of the whole network or the effective neuron is a discrete-time Bernoulli process. Denote the firing rates of excitatory and inhibitory neurons by *r*_*E*_ and *r*_*I*_, respectively, they depend on the strength *α* of *I*_noise_ and on *g*_*E*_ and *g*_*I*_ (see below).

The probability of the effective neuron having a null event in a time bin is 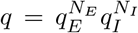, where 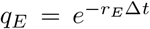 and 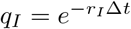. Then

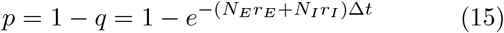

is the probability of the effective neuron having a spiking event in a time-bin. The duration *T* of an avalanche is equal to the nonzero interval between two consecutive null events of the effective neuron. Using the known result of probability distribution of inter-null-event interval for Bernoulli process [68], we obtain

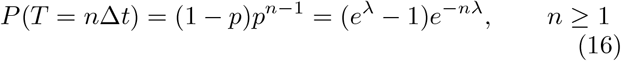

which are exponential and

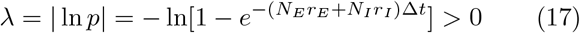

decreases with Δ*t*. The conditional size distribution of avalanches with duration of one time bin can be easily obtained:

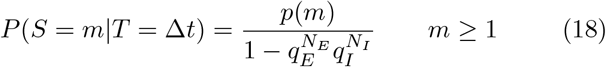

where *p*(*m*) is the probability of having *m* spikes in one bin:

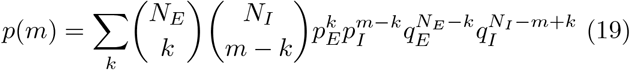

and the sum over *k* is from max{0, *m* − *N*_*I*_ } to min{*m, N*_*E*_}. Here, we make the assumption that every neuron at most spikes once within one time bin, which is justified by the low firing rates in state I. As the spiking activities in each bin are independent,

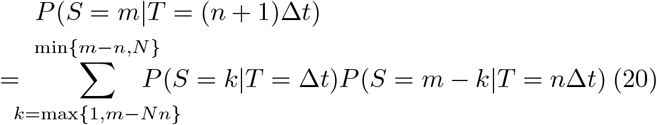

with *n* +1 ≤*m* ≤ (*n* +1)*N*. Using Eq. (18) and Eq. (20), we obtain

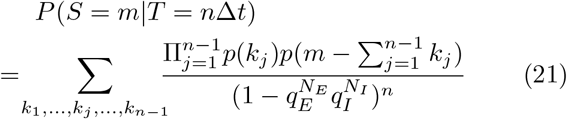

for *n* ≥2, and the sum over *k*_*j*_, *j* = 1, 2, …, *n* 1, is from max {1, *m* − (*n* −*j*)*N*} to min {*m* − (*n* −*j*), *N*}. The size distribution *P* (*S*) is thus given by

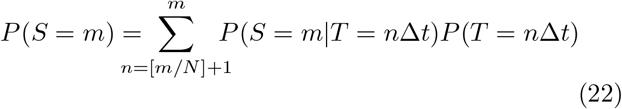

The sums in Eqs. (21) and (22) are evaluated numerically to obtain *P* (*S*), which is not an exponential function. We compare our theoretical results for *P* (*T*) and *P*(*S*), evaluated using *r*_*E*_ and *r*_*i*_ measured in the simulations, with the numerical results and excellent agreement is found (see Fig. 4). The dependence of *λ* on Δ*t* is also confirmed by the numerical results (see Fig. 8).

**FIG. 8:**
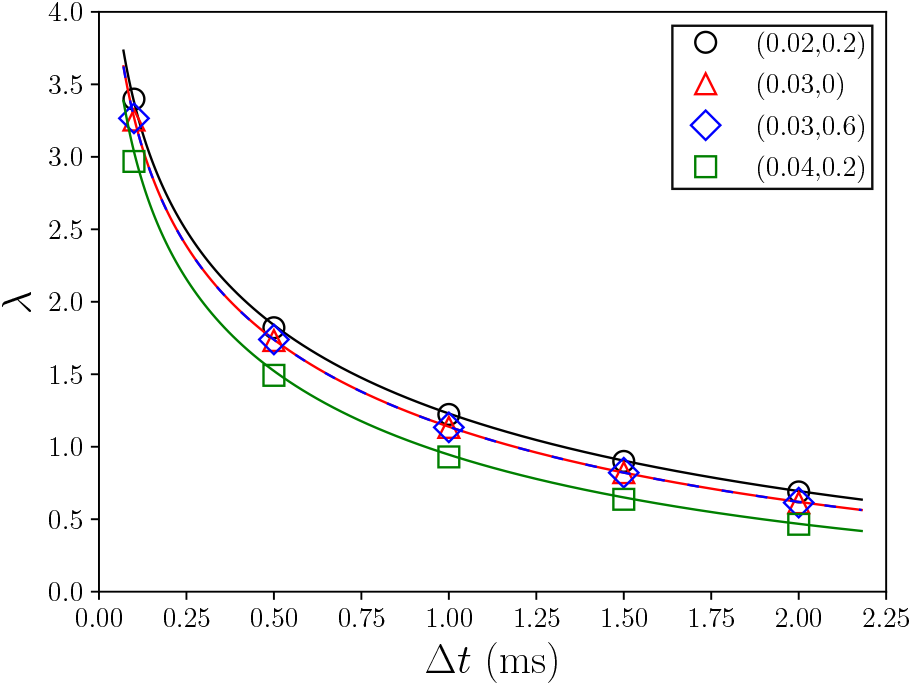
Dependence of λ for *p*(*T*) on Δ*t* for different values of (*g*_*E*_,*g*_*I*_) in state I. The theoretical result Eq. (17) (solid line) is in excellent agreement with the numerical results.

**FIG. 9:**
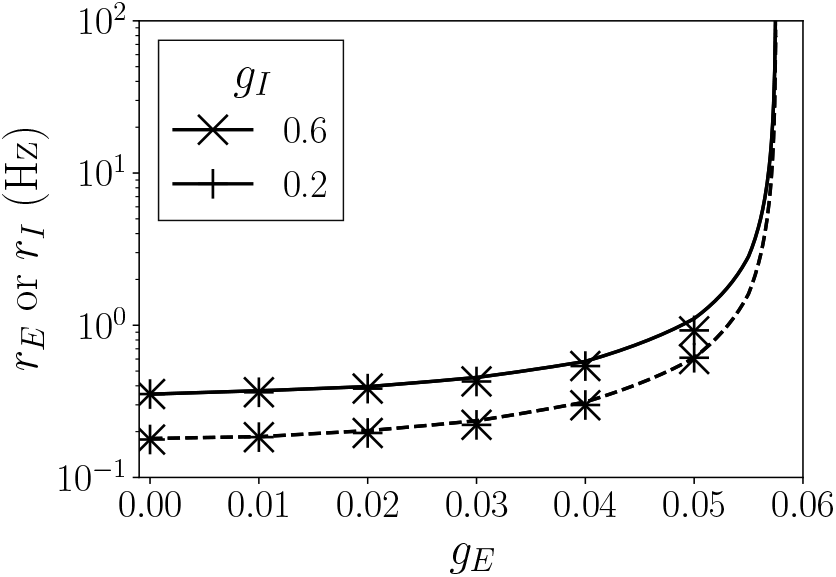
Dependence of the firing rates *rE* and *rI* on *gE* for two fixed values of *gI*. The numerical values (symbols) are in good agreement with the theoretical results (solid lines) given by Eq. (25).

Next, we study the dependence of RE and *r*_I_ on *g*_*E*_ and *g*_*i*_ and show that state I would become unstable when *g*_*E*_ is larger than some threshold value. Let the bare firing rate of an isolated excitatory or inhibitory neuron induced solely by the stochastic current *I*_noise_ be 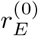 or 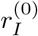 In the presence of coupling with the other neurons in the network, a single spike from a presynaptic excitatory (inhibitory) neuron of a certain neuron, would cause a jump in the excitatory (inhibitory) conductance and generate a small presynaptic current. This presynaptic current by itself cannot induce further spikes but would change the probability of spike initiation by the stochastic current in that neuron and thus change its firing rate. Denote the changes in the firing rates due to the bare firing rates as 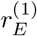 and 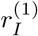. Similarly,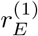 and 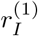 would induce additional changes in firing rates, denoted by 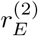 and 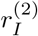, and this process continues. The changes in firing rates in two consecutive generations are related:

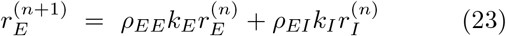

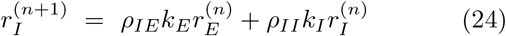

for *n* = 0,1,2,…. Here, *ρxy* is the change in firing rate of a neuron of type *X* (*E* or *I*) due to one single spike of a presynaptic neuron of type *Y* (*E* or *I*) and *k*_*E*_(*I*) is the incoming excitatory or inhibitory degree of a neuron. The actual firing rates *r*_*E*_ and *r*_*I*_ are the sums of the bare firing rates and the changes in the firing rates induced in all generations, thus

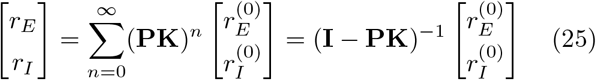

where the matrices **P** and **K** are defined by

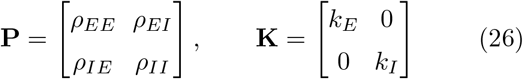

When the determinant of the matrix (**I PK**) becomes zero, the firing rates would diverge indicating that the state of independent and irregular spiking becomes unstable and a transition to a new dynamical state with correlated neuron spiking would occur. We measure *ρ*_*EE*_, *ρ*_*IE*_, *ρ*_*EI*_ and *ρ*_*II*_ by numerical simulations of isolated neurons subjected to one additional spike with *g*_*E*_ or *g*_*I*_, and verify Eq. (25) as shown in Fig. 8. By plotting the determinant of (**I**− **PK**) as a function of *g*_*E*_ for different fixed values of *g*_*I*_, we obtain the value 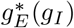 at which the determinant vanishes. As shown in Fig. 1, 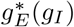 is slightly above the boundary between states I and II.

### B. State II

One prominent feature of state II is the coherent network bursting manifested in an oscillatory population averaged time-binned firing rate *r*(*t*) as shown in Fig. 3. This leads us to the idea that the collective dynamics in state II could be captured by the stochastic dynamics of the effective neuron with a time-dependent firing rate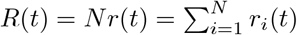, which can in turn be approximated by an inhomogeneous Poisson process with the time varying rate *R*(*t*). For small *g*_*I*_, the peaks in the oscillatory *R*(*t*) are similar whereas for larger *g*_*I*_, there are large variations in the heights, widths and shapes of the peaks in the oscillatory *R*(*t*) (see Fig. 3). Therefore, to test this idea, we focus on small *g*_*I*_ so that we can approximate *R*(*t*) as a repetition of one typical peak. To extract *R*(*t*) of one typical peak, we use a time bin of width of 1 ms, isolate the peak by truncating data on both sides when there are no spiking activity in at least 5 consecutive time bins, and smooth the data using a Gaussian filter. Then we generate spiking data for an inhomogeneous Poisson process with this time varying rate *R*(*t*) as follows. Using a small *δt* = 0.001 ms such that *R*(*t*) varies slowly in *δt*, we generate the number of spikes *k* in each time interval [*t, t* + *δt*] according to the Poisson distribution

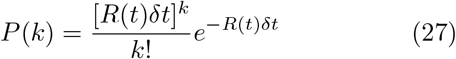

We repeat this process 4000 times to get good statistics. Using this generated time series of the number of spikes, we carry out avalanche analysis using a time bin Δ*t* equal to IEI_ave_ found in the network simulation. As shown in Fig. 10, *P* (*S*) and *P* (*T*) and ⟨*S*|*T*⟩so obtained match perfectly the numerical results found in the network simulation. This perfect matching clearly demonstrates that the power-law distributed neuronal avalanches with properties described by Eqs. (1)-(4) are direct consequences of stochasticity and the oscillatory *R*(*t*) arising from coherent bursting and criticality does not play a role. These results thus confirm earlier ideas that power-law distributed neuronal avalanches could result from stochastic dynamics [40, 41, 43] and, in addition, pinpoint the origin of the time-dependent rate [43, 45].

**FIG. 10:**
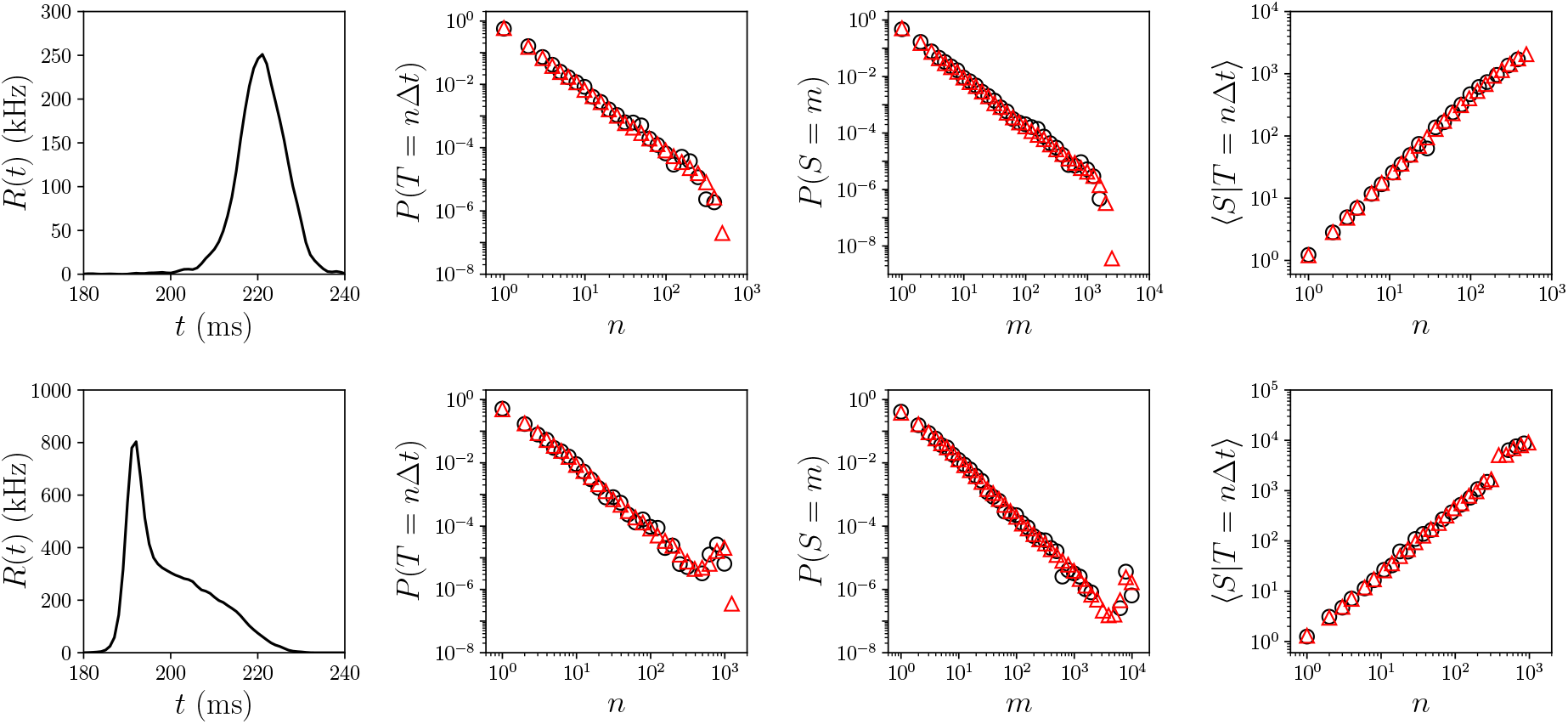
Comparison of *P*(*T*), *P*(*S*) and ⟨*S*|*T*⟩obtained from inhomogeneous Poisson process with time-dependent rate *R*(*t*) and from the neuronal network in state II for (*gE*, *gI*) = (0.1, 0.2) (top) and (*gE*, *gI*) = (0.3, 0.2) (bottom). *R*(*t*) is equal to one typical peak of the oscillatory population firing rate. Results from simulations of the inhomogeneous Poisson process and the neuronal network are shown as triangles and circles, respectively.

It has been shown that in networks of adaptive neurons, coherent network bursts result from the cooperation between strong excitation and sufficiently strong adaptation [66, 69, 70]. This suggests that the existence of state II would require the adaptation strength to be compatible with the excitatory synaptic strength. We investigate this issue using numerical simulations with varying adaption strength by replacing *d* in Eq. (8) with *κd*, and varying *κ*:

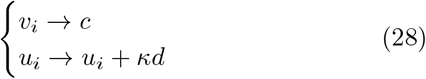

We map the phase diagram and locate the boundaries between the states in the *g*_*E*_-*κ* parameter space for a few values of *g*_*I*_ as shown in Fig. 10. The boundary between states I and II does not depend on *κ*, which is not surprising. In state I, there is no prolonged stimulus as firing activities are triggered by the stochastic input only thus the stability of state I would not be affected by the strength of spike adaptation or *κ*. The boundary between states II and III clearly depends on *κ* for fixed *g*_*E*_ and *g*_*I*_. In particular, if a network is in state II at *κ* = 1, it would undergo transition to state III as *κ* is decreased below 1 with *g*_*E*_ and *g*_*I*_ held fixed. Similarly, a network in state III at *κ* = 1 undergoes transition to state II as *κ* is increased above 1 with *g*_*E*_ and *g*_*I*_ held fixed. At a fixed value of *g*_*E*_ that is greater than 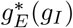, the boundary between states II and III defines the curve *κ*^*∗*^(*g*_*E*_, *g*_*I*_) *>* 0 above which state II exists and below which state III exists. It can be seen that *κ*^*∗*^ increases with *g*_*E*_ at fixed *g*_*I*_ and decreases with *g*_*I*_ at fixed *g*_*E*_. At a fixed level of adaptation with a fixed value of *κ* greater than *κ*^*∗*^, state II thus exists in a range of *g*_*E*_ given by 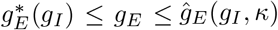, where 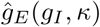 is obtained from *κ*^*∗*^(*g*_*E*_, *g*_*I*_) = *κ*. Moreover, in the absence of adaptation at *κ* = 0, the dynamics of the network undergoes transition from state I directly to state III when *g*_*E*_ exceeds 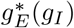 for all values of *g*_*I*_. Hence, the existence of state II and thus the power-law distributed neuronal avalanches requires an adaptation that is sufficiently strong to balance the excitation.

Finally, we show that the four observed properties, Eqs. (1)-(4), of neuronal avalanches are related if the conditional size distribution for avalanches of a given duration *P* (*S*|*T*) obey a scaling form:

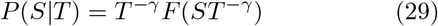

where *F* (*x*) is a scaling function. Specifically, Eq. (3) follows from Eq. (29), and Eq. (2) and Eq. (4) follows from Eq. (1) and Eq. (29) if the discrete distributions are approximated by continuous ones and 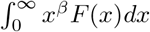 is finite for *β >* 0. First,

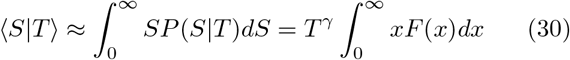

which gives Eq. (3). Second,

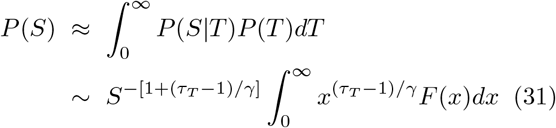

and hence, *P* (*S*) is power-law distributed, as described by Eq. (2), and *τ*_*S*_ = 1+(*τ*_*T*_ − 1)*/γ*, which is just Eq. (4). Our derivation differs from [36] which derives Eq. (4) using Eq. (1) and (2) and the assumption that *P* (*S*|*T*) is a delta function. As shown in Fig. 12, *P* (*S T*) are different from a delta function, and the curves of *P* (*S*|*T*) for different values of *T* indeed collapse approximately into one single curve when suitably rescaled. Thus, *P* (*S*|*T*) obeys the scaling form Eq. (29) approximately in both the neuronal network and the inhomogeneous Poisson process.

**FIG. 11:**
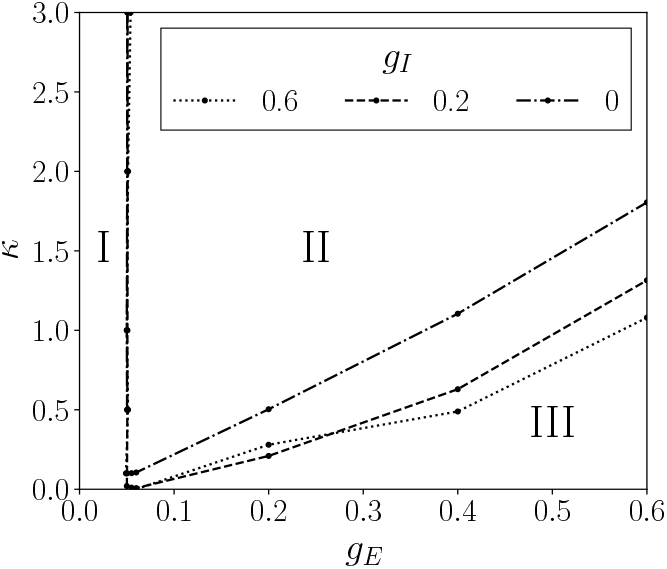
Boundaries between states I, II, and III in the *κ*-*gE* parameter space for different values of *gI*. The boundary between state II and state III is *κ*∗(*gE*, *gI*).

**FIG. 12:**
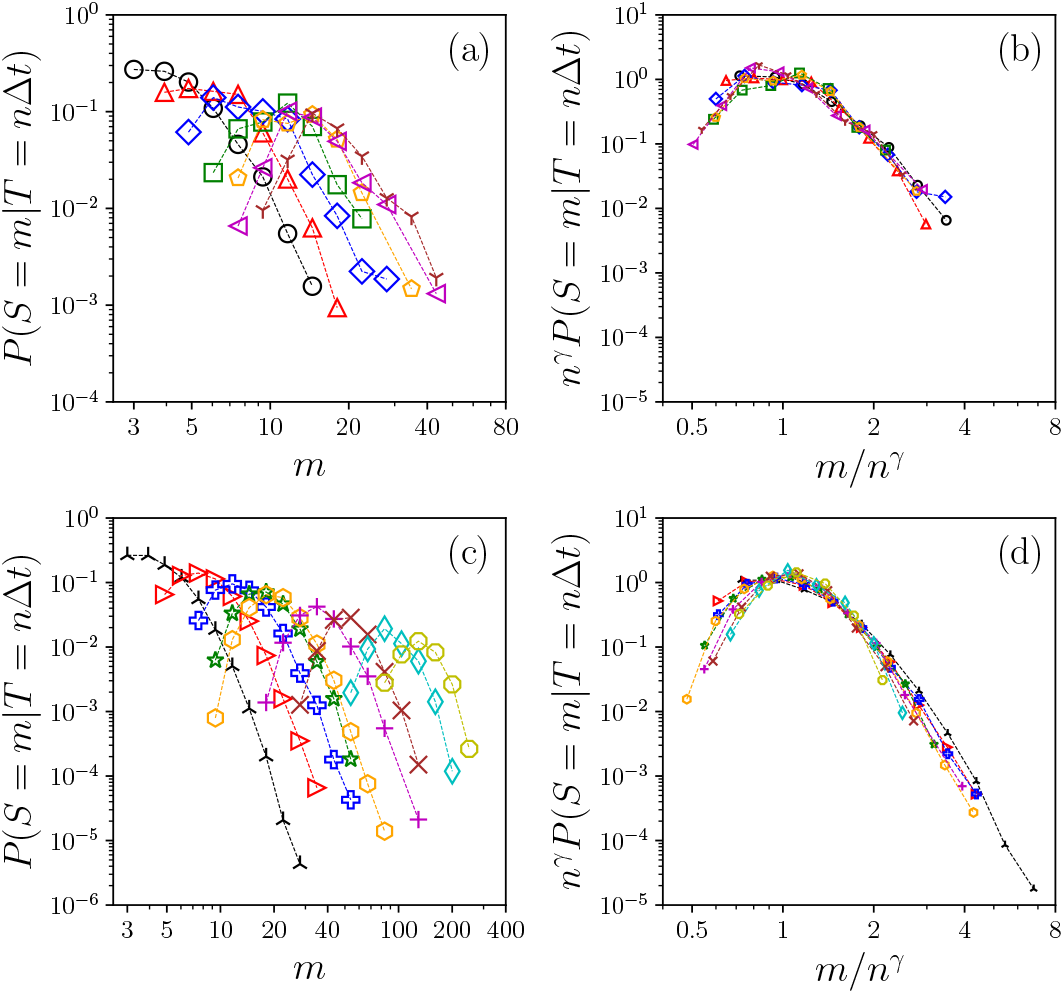
Scaling of *P*(*S = m*|*T = n*Δ*t*) for (a)-(b) neuronal network with *n* = 3-9 and (c)-(d) inhomogeneous Poisson process with *n* ranging from 3 to 40 for (*gE*, *gI*) = (0.3, 0.2) in state II. *γ* = 1.3 for (b) and 1.29 for (d).

### C. State III

When *g*_*E*_ is larger than 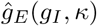, the adaptation is too weak to maintain the coherent bursting. The system changes to state III in which the time-binned population firing rate exhibits only weak fluctuations around the average firing rate *R*_0_ of the whole network. Here, *R*_0_ is the total number of spikes of all the neurons divided by the total observation time. Therefore, the dynamics of state III can be approximated by a standard Poisson process with a constant rate *R*_0_. For constant *R*_0_, analytical results for *P* (*T*) and *P* (*S*) have been derived [45]:

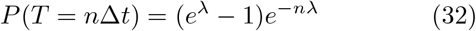

and

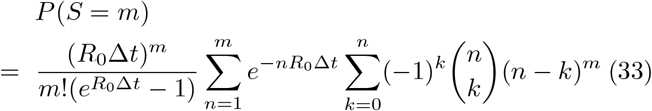

where 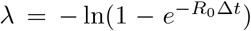 Thus, *P* (*T*) is exponential as in state I and indeed Eq. (16) can be written as Eq. (32) since *N*_*E*_*r*_*E*_ + *N*_*I*_ *r*_*I*_ = *R*_0_ in state I. However, *R*_0_ ≠ *N*_*E*_*r*_*E*_ + *N*_*I*_ *r*_*I*_ in state III as the firing rates differ among the excitatory and inhibitory neurons. Moreover, the dynamics of the neurons in state III are not independent and cannot be approximated as *N* independent Poisson processes. The distributions *P* (*S*) are close to exponential similar to state I. These analytical results are in good agreement with our numerical results, as shown in Fig. 4.

## V. DISCUSSIONS AND CONCLUSIONS

Spontaneous brain activity displays rich spatial and temporal dynamical structures such as oscillations and power-law neuronal avalanches. Understanding its origin, underlying mechanism and functional significance is a fundamental challenge in neuroscience. The observed power-law distributed neuronal avalanches have been interpreted as evidence supporting the hypothesis that the brain is operating at the edge of a continuous phase transition. However, this critical brain hypothesis remains controversial. Moreover, there is not yet a study showing oscillations and power-law neuronal avalanches with the observed features, Eqs. (1)-(4) and (*τ*_*T*_−1)*/*(*τ*_*S*_−1) ≈1.3, using a model with biological details.

In this paper, we have shown that in networks of adaptive neurons subjected to stochastic input, with biophysically plausible Izhikevich’s model of neurons and conductance-based synapses, a dynamical state emerges that displays the above mentioned features of spontaneous activity. We have mapped out the dynamics for the simple network A with homogeneous excitatory and inhibitory incoming degrees and constant excitatory and inhibitory synaptic strengths *g*_*E*_ and *g*_*I*_ and found three distinct dynamical states: states I, II and III. State II exists for a range of intermediate values of *g*_*E*_ and the dynamics in state II have features resembling those of spontaneous activity observed in experiments. Similar results have been found in two additional networks, one network with the same network structure but with a uniform distribution of excitatory and inhibitory synaptic strengths centered at *g*_*E*_ and *g*_*I*_ (network B) and the other network with heterogenous incoming degrees and long-tailed outgoing degree distribution (network C).

The system is in state II when there is sufficiently strong excitation with *g*_*E*_ exceeding 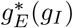 and when the strong excitation is balanced by adaption, with *g*_*E*_ below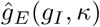, where *κ* measures the strength of adaptation. At fixed *g*_*I*_, 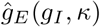 and thus the range of state II increases with *κ*. The balance of excitation and adaptation generates coherent network bursting [66, 69, 70], which is manifested as oscillations in the time-binned population firing rate *R*(*t*) of the whole network. At shorter time scales, the bursting activities give rise to avalanches of diverse sizes and durations with power-law *P* (*T*) and *P* (*S*) and features as observed in experiments, namely Eqs. (1)- (4) and (*τ*_*T*_−1)*/*(*τ*_*S*_−1) ≈1.3. We have demonstrated that these neuronal avalanches are direct consequences of stochasticity and the oscillatory *R*(*t*). We have further found the interesting result that the conditional size distribution of avalanches with a given duration, *P* (*S*|*T*), obeys approximately a scaling form Eq. (29) and shown that this result together with a power-law duration distribution, Eq. (1), can account for a power-law size distribution, the power-law dependence on *T* of the average size of avalanches with given duration *T*, and the approximate scaling relation between the exponents. Neuronal avalanches with power-law distributions in size and duration have been found in stochastic dynamics of networks of excitatory and inhibitory integrate-and-fire neurons without adaptation, using the model by Brunel [65], but it was found that the scaling relation Eq. (4) was not satisfied [43]. Power-law distributed avalanches have also been reported in stochastic dynamics of networks of Izhikevich neurons with and without synaptic plasticity and were associated with critical behavior [7, 71]. In contrast, our work shows that criticality does not play a role. It remains to understand the conditions on *R*(*t*) for generating power-law *P* (*T*) and *P* (*S*|*T*) that obeys the scaling form Eq. (29).

When *g*_*E*_ is too small such that the synaptic current *I*^syn^ cannot excite a spike on its own and spiking of neurons are triggered by the stochastic current *I*_noise_, state I occurs. State I is a state of random and independent firing of *N* neurons, which can be well approximated by *N* independent Poisson processes. Using this approximation, we have derived analytical expressions showing exponential *P* (*T*) and close-to-exponential *P* (*S*) and these theoretical results are in excellent agreement with the numerical results found in state I. We have further shown that state I is unstable when *g*_*E*_ is greater than 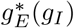 and the system would undergo transition to a state of correlated neuron spiking. When *g*_*E*_ further exceeds 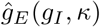, the adaptation is too weak to balance the excitation and in this state III, the spiking activities of the neurons are fast and incoherent, and the sum of these incoherent spiking activities is thus averaged out to give a population firing rate that is approximately constant in time. The collective dynamics of state III can then be approximated by a standard Poisson process of constant rate and the duration distribution of the neuronal avalanches are exponential [45] as in state I and the size distributions are close to exponential [45] similar to state I.

In conclusion, we have shown that the collective stochastic dynamics of adaptive neurons display oscillations, arising from coherent bursting, as well as power-law neuronal avalanches with features as observed in spontaneous activity in the entire state II. These results suggest that the brain is likely operating in this state II. Our work has shown that oscillations and power-law neuronal avalanches do not just coexist in state II but the power-law neuronal avalanches are direct consequences of stochasticity and the oscillations, coherent bursting is itself the result of the balance between excitation and adaptation, and criticality does not play a role. Moreover, inhibition or excitation-inhibition balance, hierarchical network structure or heavy-tailed synaptic strength distribution are not necessary factors for power-law neuronal avalanches, contrary to suggestions made in earlier studies [22–24, 26].

## Appendix Results for Networks B and C

Network B is modified from network A with the excitatory and inhibitory synaptic strength taken from uniform distributions with mean *g*_*E*_ and *g*_*I*_ and width Δ = 0.04. Network C is reconstructed from multi-electrode array data measured from a neuronal culture [62] using the method of directed network reconstruction proposed by Ching and Tam [63] and with the excitatory and inhibitory synaptic strengths taken to be constant values *g*_*E*_ and *g*_*I*_. In network C, *N* = 4095 and the distributions of excitatory and inhibitory incoming degrees are bimodal while the distributions of outgoing degree are long-tailed, which are qualitatively different from those of network A, as shown in Fig. 13.

**FIG. 13:**
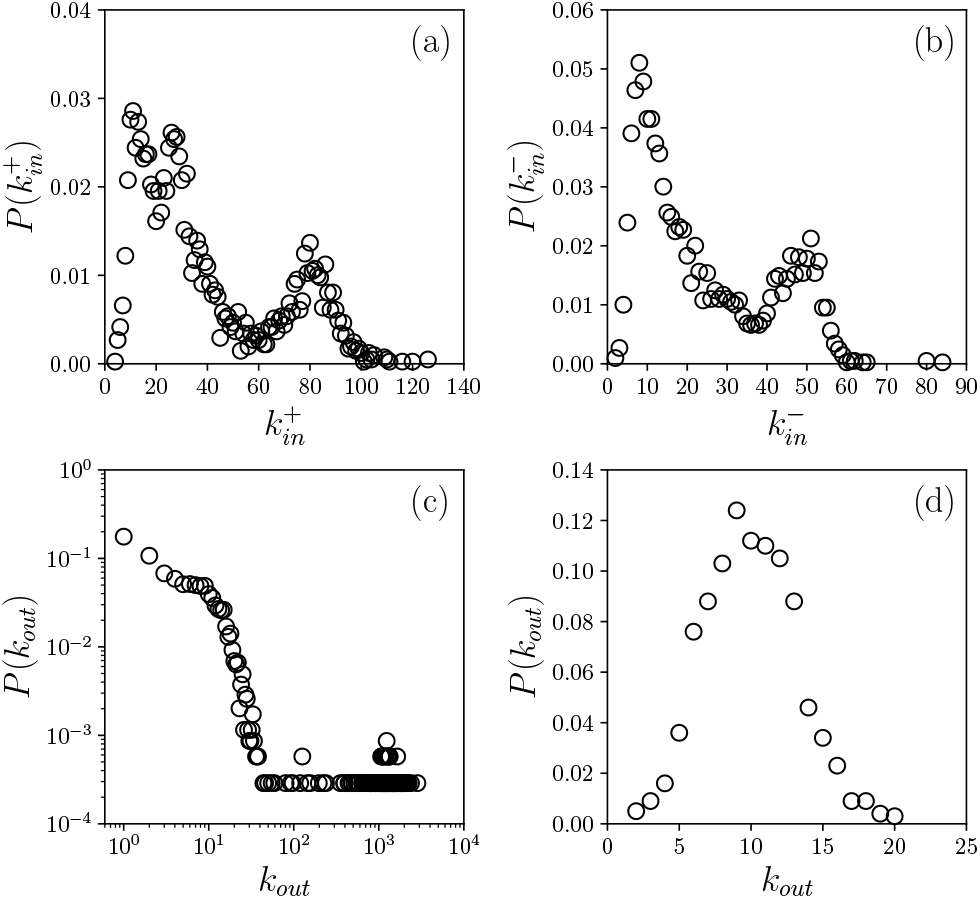
Distributions of incoming and outgoing degrees of network C. (a): Distribution of the excitatory incoming degree 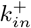, (b) Distribution of the inhibitory incoming degree 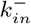 and (c) Distribution of the outgoing degree *kout*. For comparison, we show the distribution of *kout* of network A in (d).

We have integrated the stochastic differential equations (5) and (6) together with equations (9) and (10) using the Euler-Maruyama method [64] with a time step *dt* = 0.02 ms for network B and *dt* = 0.001 ms for network C to study the dynamics as a function of *g*_*E*_ and *g*_*I*_ at fixed values of *α*. Similar to network A, three dynamical states have been found for networks B and C. A similar phase diagram has been mapped out for network B at *α* = 5 (see Fig. 14). For network C, we focus to study the statistics of the neuronal avalanches found in state II at *α* = 3. As in network A, the distributions *P* (*T*) and *P* (*S*) are well described by power law for a range of of values of Δ*t* and we take Δ*t* = IEI_ave_ or a value around the middle of the range when power-law is not observed at Δ*t* = IEI_ave_. We present the results for the exponents for all the cases studied for network C in Table 2. The exponents again obey the scaling relation Eq. (4) approximately with (*τ*_*T*_−1)*/*(*τ*_*S*_−1) ≈1.3 (see Fig.15.).

**TABLE 2:** Exponents τ_*T*_, τ_*S*_ and γ found for different (*g*_*E*_, *g*_*I*_) in state II of network C. When Δ*t*≠ IEI_ave_, the values of IEI_ave_ are indicated in parentheses.

| $g_E$ | $g_I$ | $\tau_T$ | $\tau_S$ | $\frac{(\tau_T - 1)}{(\tau_S - 1)}$ | $\gamma$ | $\Delta t$ (ms) |
| --- | --- | --- | --- | --- | --- | --- |
| 0.03 | 0.06 | 2.37 | 2.07 | 1.28 | 1.29 | 0.02 (0.11) |
| 0.0375 | 0.06 | 2.37 | 2.07 | 1.28 | 1.28 | 0.015 (0.075) |
| 0.045 | 0.06 | 2.26 | 1.95 | 1.33 | 1.29 | 0.055 |
| 0.0525 | 0.06 | 2.14 | 1.88 | 1.30 | 1.30 | 0.04 |
| 0.06 | 0.06 | 2.22 | 1.94 | 1.30 | 1.32 | 0.035 |
| 0.0675 | 0.06 | 2.19 | 1.90 | 1.32 | 1.27 | 0.03 |
| 0.075 | 0.06 | 2.31 | 2.02 | 1.28 | 1.25 | 0.01 (0.03) |
| 0.0975 | 0.06 | 1.95 | 1.76 | 1.25 | 1.31 | 0.025 (0.02) |
| 0.12 | 0.06 | 1.92 | 1.71 | 1.30 | 1.29 | 0.02 |
| 0.135 | 0.06 | 1.92 | 1.70 | 1.31 | 1.27 | 0.015 |
| 0.06 | 0.015 | 2.29 | 2.00 | 1.29 | 1.29 | 0.04 |
| 0.06 | 0.03 | 2.27 | 1.99 | 1.28 | 1.28 | 0.04 |
| 0.06 | 0.045 | 2.26 | 1.96 | 1.31 | 1.29 | 0.005 (0.035) |
| 0.06 | 0.075 | 2.10 | 1.86 | 1.28 | 1.26 | 0.02 (0.035) |
| 0.06 | 0.09 | 2.19 | 1.92 | 1.29 | 1.28 | 0.025 (0.035) |
| 0.06 | 0.105 | 2.22 | 1.94 | 1.30 | 1.26 | 0.02 (0.04) |
| 0.06 | 0.12 | 2.27 | 1.96 | 1.32 | 1.27 | 0.03 (0.04) |

**FIG. 14:**
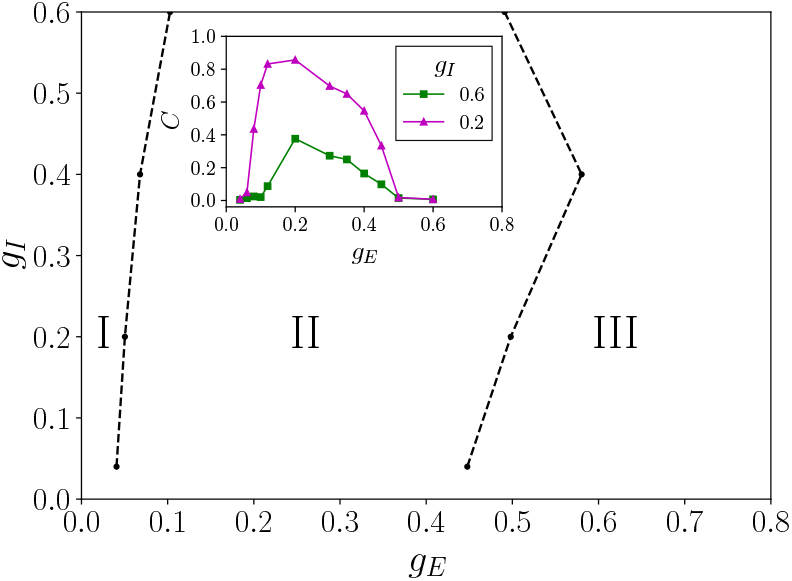
Different dynamical states in the *gE*-*gI* parameter space at *∝* = 5 for network B. The inset shows the dependence of the coherence parameter *C* as a function of *gE* at fixed values of *gI*.

**FIG. 15:**
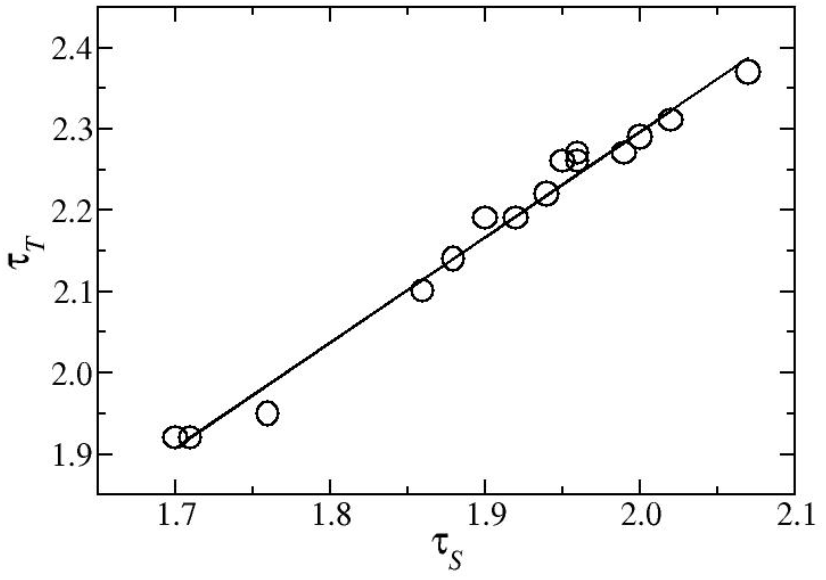
Relation between τ_*T*_ and τ_*S*_ in state II of network C for all the cases studied as shown in Table 2. The data are fitted by a straight line (*τ*_*T*_ ™1)/(*τ*_*S*_ ™1) = *K* solid line) and the fitted value of K is 1.3 for network C.

